# Brain states govern the spatio-temporal dynamics of resting state functional connectivity

**DOI:** 10.1101/832998

**Authors:** Felipe Aedo-Jury, Miriam Schwalm, Lara Hamzehpour, Albrecht Stroh

## Abstract

Previously, using simultaneous task-free fMRI and optic-fiber-based neuronal calcium recordings in the anesthetized rat, we identified BOLD responses directly related to slow calcium waves, revealing a cortex-wide and spatially organized BOLD correlate (Schwalm et al. 2017). Here, with these bimodal recordings, we reveal two distinct brain states: persistent state, in which compartmentalized network activity was observed, including defined subsets such as the default mode network; and slow wave state, dominated by a cortex-wide component, suggesting a strong functional coupling of brain activity. In slow wave state we find a correlation between slow wave events and the strength of functional connectivity. These findings suggest that indeed down-up transitions of neuronal excitability drive cortex-wide functional connectivity. This study provides strong evidence that previously reported changes in functional connectivity are highly dependent on the brain’s current state and directly linked with cortical excitability and slow waves generation.

## Introduction

The spatio-temporal dynamics of cortical excitability are constantly impacted by fluctuations of internally generated activity, even in the absence of external inputs. The task-free state of the cortical functional architecture is characterized by canonical “*resting state networks*”^1–3^. Specific resting state networks have been identified in humans ^4^, primates ^5^ and rodents ^6,7^ using task-free fMRI methods. However, the spatio-temporal features of these networks are not static ^8^. Alternating states of excitability - or brain states – can dynamically occur, even in different regions at the same time, as seen in the awake condition^9^. Brain states consist in internally generated activity and influence responses upon incoming sensory afferents^10,11^. Indeed, brain state changes are accompanied by ostensive alterations in the spatiotemporal pattern of neuronal activity in many brain areas, observed in local electrophysiological^7,8,12^ and optical^6,13^ recordings, as well as large-scale readouts of brain activity, such as BOLD fMRI^14–16^. This calls for a brain-state informed assessment of resting state functional connectivity^17^.

Recently, we have identified two brain states which were shown to occur under different types of anesthesia used for small animal fMRI studies^11^. In persistent state, which can occur during but is not limited to awake periods, neurons are rather depolarized, sparsely active, leading to temporally dynamic, modality specific, network configurations^18^. In contrast to the persistent state, stands the bimodal activity pattern of slow oscillations, or slow wave state which has been extensively described ^5,11,19–25^, but only most recently in the framework of BOLD fMRI^11,26,27^. The corresponding low-frequency component ranges at 0.2-1 Hz, reflecting alternating activity patterns: active (*“up”*) periods in which cells are depolarized and fire action potentials in temporally restricted periods, and silence (*“down”*) periods with rather hyperpolarized membrane potentials and an almost complete absence of neuronal activity ^28,29^. Slow waves can occur rather locally^30^ or spread over the entire cortex^11,19^, allowing activity to propagate^21^. In this work we found that independent component analysis in persistent state revealed a canonical resting state associated networks, such as the default mode^11,31^ or sensory networks^32,33^. The concept of brain states as defined by the local and global features of network dynamics by no means entails, that different behavioral states as e.g. anesthetized condition or natural sleep would be identical, however, defining features of the slow wave state, such as bimodality, can be identified in a plethora of different conditions^19,24,34,35^ as the fundamental difference between fast and slow timescale cortical dynamics is likely regulated by distinct mechanisms at the single neuron level^36^.

Task-free fMRI functional connectivity analysis is used to identify and characterize neural networks in health and disease^37,38^, during learning^15^ or memory consolidation^16^. The possibility to combine task-free fMRI with invasive recordings of brain activity constitutes a pivotal branch of translational research^39^. Surprisingly, in rodents as in humans, little has been investigated on how the brain’s current state impacts functional connectivity derived from BOLD activity, despite evidence that, for instance, small changes in anesthesia can dramatically change brain connectivity^40,41^. While variations in functional connectivity have been related to cognitive state^42^ in humans, the effect of neurophysiologically defined states on functional connectivity in the cortex remains unknown. Directly relating functional connectivity to defined brain states may have explanatory value for a broad range of functional brain connectivity studies. We propose that connectivity measures in different conditions such as different anaesthetics ^14,43–46^ can be classified based on the respective brain state. Based on these classifications, meaningful comparisons can be drawn both between functional connectivity signature in rodents and humans.

Here, we find distinct BOLD functional connectivity patterns for persistent state and slow wave state, revealing compartmentalized network activity during persistent state but a cortex-wide component during slow wave state. We furthermore, unveil a massive cortex-wide functional coupling of brain activity during slow wave state and identified its origin in the transitions from population *down* to population *up* states occurring during the generation of the slow waves. Our data reveals the imminent consequences in terms of functional networks configuration of each slow wave event.

## Materials and Methods

### Animals

Experiments were performed on 17 adult female Lewis rats (> 12 weeks old, 160-200 g), of which 8 were used for fMRI measurements, 5 were implanted with an optic fiber in the visual cortex for combined optic fiber-based calcium recordings/fMRI measurements and 4 were used for optical recordings on the bench. Animals were housed under a 12 h light–dark cycle and provided with food and water ad libitum. Animal husbandry and experimental manipulation were carried out according to animal welfare guidelines of the Johannes Gutenberg-University Mainz and were approved by the Landesuntersuchungsamt Rheinland-Pfalz, Koblenz, Germany.

### fMRI Data acquisition

fMRI data acquisition for connectivity analyses was performed in two sessions for each animal. For the first session, animals were anesthetized with isoflurane (1.2-1.8 % in 80 % air and 20 % oxygen). For the second session animals were sedated with meditomedine (Domitor, bolus injection of 0.04 mg/kg i.p. followed by continuous i.p. infusion of 0.08 mg/kg/hr). The order of the sessions was randomly assigned to each animal. Both sessions were separated by at least one week. Measurements were performed on a 9.4 T small animal imaging system with a 0.7 T/m gradient system (Biospec 94/20, Bruker Biospin GmbH, Ettlingen, Germany). Each experimental protocol consisted of T1-weighted anatomical imaging (image size 175 × 175 × 80, FoV 26 × 29 × 28 mm, resolution 0.15 × 0.165 × 0.35 mm), 30 minutes of resting state T2*-weighted images acquired with a single-shot gradient echo EPI sequence (TR = 1.5 s, TE = 14 ms and FA 65°, 320 × 290 μm^2^ spatial resolution, slice thickness 0.8 mm, 34 continuous slices, 1200 acquisitions) and a high resolution in-plane T2-MSME images of the 34 slices (TE = 12 ms, 100 × 100 μm^2^ spatial resolution, image size 256 × 256, slice thickness 0.8 mm, 34 continuous slices). The same protocol was used in the combined recordings with optic-fiber based calcium recordings. Body temperature and breathing rate of the animal were monitored and recorded during the entire experimental procedure using a SA instruments small animal MRI-compatible monitoring system (Model 1030, Small Animal Instruments, Inc., Stony Brook, NY, USA).

### fMRI analysis

#### Data preprocessing

GLM fMRI data processing was performed using Brain Voyager 20.6 for windows (Brain Innovation, Maastricht, Netherlands). Preprocessing steps included slice scan time correction, 3D motion correction and smoothing of the images, using cubic-spline interpolation method and trilinear / sinc interpolation for the 3D motion correction. Datasets were smoothed using a 0.8 mm FWHM Gaussian Kernel. The initial 5 images were discarded to include only signals that have reached steady state. T1 images were manually aligned to the in vivo MRI template of Valdes-Hernandez et al. ^47^ and the T2*-weighted images were manually aligned to the T1 images already aligned to the MRI template used as for the high resolution in-plane T2*-weighted images. The same preprocessing was performed for the recordings combined with optic-fiber-based calcium recordings.

#### Independent component analysis

For the independent component analysis the freely available software “Group ICA of fMRI Toolbox (GIFT)”, version 4.0b (http://mialab.mrn.org/software/gift/; RRID:SCR_001953) was used. Mask-calculation using the first image was employed and time-series were linearly detrended and converted to z-scores. For back-reconstruction, the “GICA”-algorithm ^48^ has been used and all results have been scaled by their z-scores. We used the Infomax-algorithm with regular stability analysis without autofill. Images for the ICA were motion corrected and smoothed as described above for each animal. For visualization purposes, the z-scores of each animal were averaged and then plotted in Brain Voyager over the template (Figure 1).

**Figure 1:**
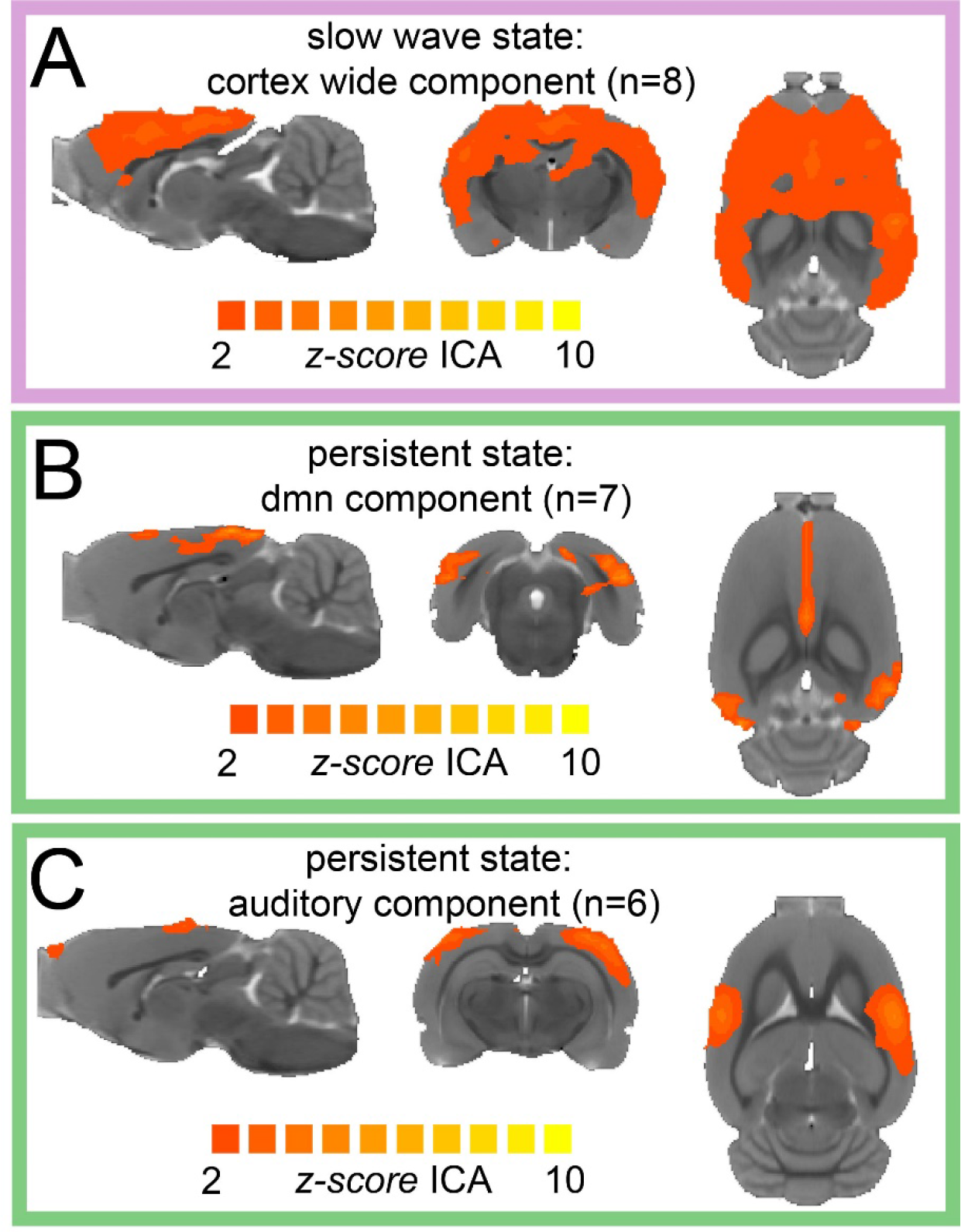
ICA reveals distinct components for each brain state. A- Average z-score maps of cortex wide component during slow wave state. B-C Average z-score maps of two components found in 7 and 6 animals respectively (B default mode network, C auditory component) during persistent state.

#### Connectivity analysis

Connectivity analyses were performed using in-house built Matlab scripts (The Mathworks, Inc., Natick, MA, USA). BOLD-signal was first normalized to its mean intensity. Then, cortical volume was parcellated in 96 regions of interest (ROIs) using the MRI template of Valdes-Hernandez *et al* ^47^ and the average signal for each area was calculated. The average signal of the ventricles and white matter was extracted and together with the breathing signal added as a covariable. In a second connectivity analysis, the time course of the cortex-wide component was added as a covariable, either in its original form or temporally reversed as a control. Then, partial correlation coefficients were calculated for each pair of ROIs. The significant values were calculated for each matrix using the r-value (0.348±0.045 s.e.m) obtained after false discovery rate (FDR) correction.

#### fALFF analysis

The amplitude of low-frequency fluctuations (ALFF) of the task free fMRI signal is the power of the signal within a certain frequency range and measures the intensity of regional spontaneous brain activity in humans^49^ as a measure of local metabolism in a particular region. To reduce sensitivity to physiological noise, Zou et al.^50^ introduced fractional ALFF (fALFF), which is defined as the ratio of low-frequency amplitudes (0.01–0.08 Hz) to the amplitudes of the entire frequency range (0–0.25 Hz). Similarly, we conducted fALFF analysis using the ratio of 0.01-0.1 Hz to the entire frequency range 0-0.25 Hz. We calculated the fFALFF value for each voxel and then averaged those values for each ROI and compared the distribution of the 96 ROIs for both brain states (slow wave state and persistent state).

#### Euclidean distance

The correlation between r-scores and the Euclidean distances between the ROIs were calculated between each pair of ROIs taking the average coordinates of the voxels belonging to each ROI as its centroid.

#### Graph theory analysis

Adjacency matrices were constructed starting from correlation matrices of the functional connectivity analysis, and after thresholding (threshold was given for each matrix based on the FDR correction) were configured as *Weighted Undirected Networks*. The network measures calculated were all derived from the adaptation of the Complex Networks Analysis toolbox *Functional Brain Connectivity* for Matlab^51^. Network *nodes* in our case were ROIs, and networks *links* were the magnitudes of temporal correlations between ROIs, obtained by correlation coefficients calculated previously.

#### Matrix similarity analysis

Matrix similarity was calculated using the Frobenius Norm of the eigenvalues of each matrix. This calculation gives a measure of the distance between the two vectors resulting from the eigenvalues of the matrix. The closer to zero the obtained norm, the closer are the vectors and therefore, the more similar are the matrices.

### Indicator injections and optic fiber placement

For surgical procedures including staining with fluorescent calcium indicator and optic fiber implantation, rats were anesthetized with isoflurane (Forene, Abbott, Wiesbaden, Germany), placed on a warming pad (37° C), and fixed in a stereotactic frame with ear and bite bars.

The skull was exposed, dried from blood and fluids, and leveled for precise stereotactic injections. Under a dissection microscope a small craniotomy was performed with a dental drill (Ultimate XL-F, NSK, Trier Germany, and VS1/4HP/005, Meisinger, Neuss, Germany). The fluorescent calcium sensitive dye Oregon Green 488 BAPTA-1 (OGB-1, Invitrogen, Life Technologies, Carlsbad, CA, USA) was prepared as described previously^52^, filtered, and injected into primary visual cortex (V1; −5.5 mm AP, +3.8 mm ML, −0.5, −0.7 and 0.9 mm DV) in the experiments combining fMRI and optic fiber-based calcium recordings, and a second craniotomy and injection were performed in the primary somatosensory cortex front limb area (S1FL; 0 mm AP, +3.5 mm ML, −0.5, −0.7 and 0.9 mm DV;) for the bench experiments including optical recordings in the two locations. After injection of the indicator, the pipette was held in place for 2 min before slowly retracting it from the tissue. After removing the cladding from the tip, an optic fiber with a diameter of 200 μm was inserted perpendicular to the dura above the OGB-1 stained area, typically at 300 μm and fixed to the skull with UV glue (Polytec, PT GmbH, Waldbrunn, Germany). A detailed procedure can be found in Stroh (Ed) ^24^ and Schwalm *et al^11^*.

### Optic-fiber-based calcium recordings

For optic-fiber based calcium recordings, custom-built setups were used (FOM and FOMII, npi Electronic Instruments, Tamm, Germany), as reported previously in Stroh (Ed) ^39^ and Schwalm *et al* ^11^. The light for excitation of the calcium indicator was delivered by a 650 mW LED with a nominal peak wavelength of 470 nm. LED power was controlled by an adjustable current source. The light beam was focused by means of a fixed focus collimator into one end of a multimode fiber which was connected to the system by an SMA connector. The recorded fluorescence signals were digitized with a sampling frequency of 2 kHz using a multifunction I/O data acquisition interface (Power1401, Cambridge Electronic Design, Cambridge, UK) and its corresponding acquisition software (Spike2). Data was read into Matlab (The Mathworks, Inc., Natick, MA, USA) and downsampled from 2 kHz to 1 kHz by averaging two adjacent values. The baseline of the fluorescence signal was determined and corrected by using the Matlab function *msbackadj* estimating the baseline within multiple shifted windows of 2500 datapoints width, regressing varying baseline values to the window’s datapoints using a spline approximation, then adjusting the baseline in the peak range of the fluorescent signal. This method provides a correction for low-frequency drifts while avoiding contributions of signal-of-interest events. We employed a procedure based on exponential moving average (EMA) filters, originally established to identify slow waves in electrophysiological recordings ^23^. Onsets of slow calcium waves were defined as signal timepoints exceeding the threshold (70 %), termination of calcium slow waves was defined as signal timepoints dropping below 50 % of the threshold value. Further, the detected activity was post-processed in the following order: 1) calcium waves separated by a time-interval below 500 ms were interpreted as one wave; 2) calcium waves with a duration of less than 300 ms were discarded; 3) activity not reaching 90% of the histogram-based cumulative signal intensity was discarded. A detailed procedure can be found in Stroh (Ed) ^39^ and Schwalm *et al* ^11^.

## Results

We first conducted independent component (IC) analyses of rat task-free fMRI signals in the isoflurane-induced condition, and in the medetomidine-induced condition. We found, in the isoflurane-induced condition only, a characteristic cortex-wide component (CWC), indicative that the animal had been in slow wave state, as characterized previously^11^. Conversely, in the medetomidine-induced condition, the cortex-wide component is absent, and canonical independent components of well compartmentalized default mode networks were present, indicative of the persistent brain states. While during slow wave state we found a CWC in all animals (**Figure 1A**), during persistent state the identified components resembled those described by others in awake ^45,53^ or sedated rodents^11^. We identified default mode network activation in 7 of 8 animals (**Figure 1B**) and a bilateral somatosensory component in 6 of 8 animals (**Figure 1C**).

### Connectivity analysis

To assess whether these two IC patterns are related to different cortical connectivity patterns, we interrogated the resting state recordings to obtain the functional connectivity signature of the cortical networks during slow wave state and persistent state. We parcellated the brain based on anatomical landmarks using the Valdes-Hernandez *et al* template for rats^47^ and correlated the BOLD signal of 96 cortical structures using the average signal from the white matter structures, ventricles and breathing rate in both conditions, persistent state and slow wave state as co-variables (**Figure 2A**). We first plotted the cumulative distribution function (CDF) for the r-values of each matrix (n=8) to establish whether a difference in the number and strength of significant correlations can be found. The cutoff was calculated for each matrix using FDR (mean 0.348±0.045 s.e.m.). At the mean of the cutoff the CDF (**Figure 2B**) values differed significantly between both states (t(7)=5.223, p<=0.001). Furthermore, the number of significantly correlated pairs for each individual at its particular cutoff was significant (t(7)=6.222, p<=0.001) (**Figure 2C**). These results strongly suggest a different connectivity pattern for slow wave state and persistent state respectively.

**Figure 2:**
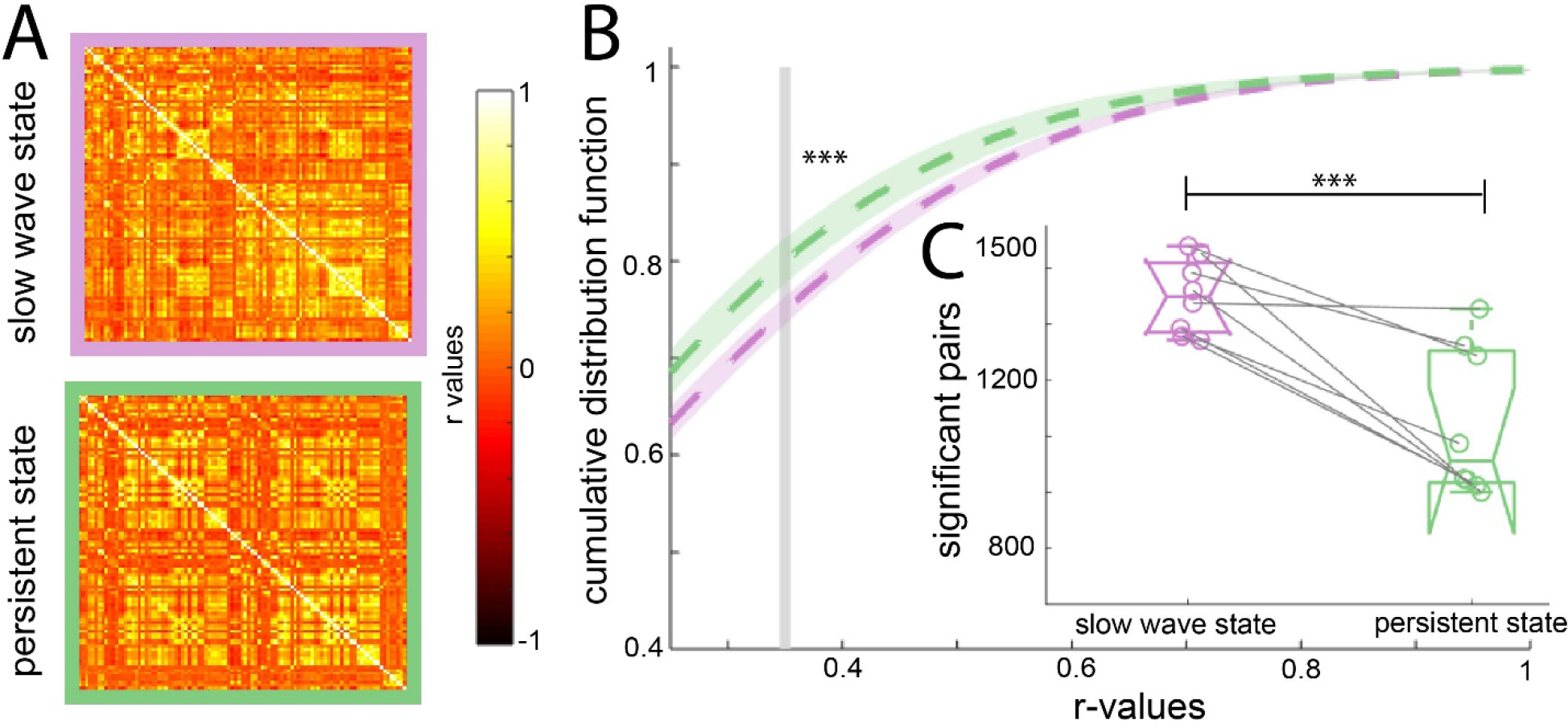
Increased connectivity in slow wave state compared to persistent state. A- Average of the correlation matrices (n=8) of the mean BOLD signal in 96 cortical regions for slow wave state(magenta) and persistent state (green). B- Cumulative distribution curve for the r-values of the correlations. The vertical gray line points the mean of the FDR values used as cutoff to identify significant correlations, ***p<0.001. C- Plot of the amount of significant correlations for each individual in both brain states ***p<0.001.

Networks during slow wave state are characterized by long periods of quiescence followed by short lapses of activity corresponding to a propagating wave^24^. fALFF analysis allows to determine the fractional power of the BOLD signal for a particular region of interest (ROI)^50^. To rule out that the connectivity values observed during slow wave state are due to the spurious consequence of an increase in power, we performed a fALFF analysis to investigate the differences in the BOLD power (**Figure 3A)**, showing that during slow wave state there is a significant shift to lower power in the distribution of the fALFF curves created with the 96 cortical ROIs of each animal (t(7)=9.65, p<=0.001). To test the dependence of the strength in connectivity from the distance of cortical areas, we correlated the Euclidean distance between pairs with the significant r-scores of the correlation of those pairs (**Figure 3B**). The closer to −90 degrees angle that the slope of those correlations gets, the more dependent of the Euclidean distance their correlation becomes. We found, that this is the case for slow wave state recordings (**Figure 3C**), whose slopes are significantly lower than those of persistent state (t(7)=5.46, p<=0.001). Altogether, these results corroborate that the BOLD signal during slow wave state is governed by different dynamics than during persistent state. Furthermore, during slow wave state, BOLD-driven connectivity seems to be less compartmentalized and shows a stronger dependency on the distance between areas, than during persistent state.

**Figure 3:**
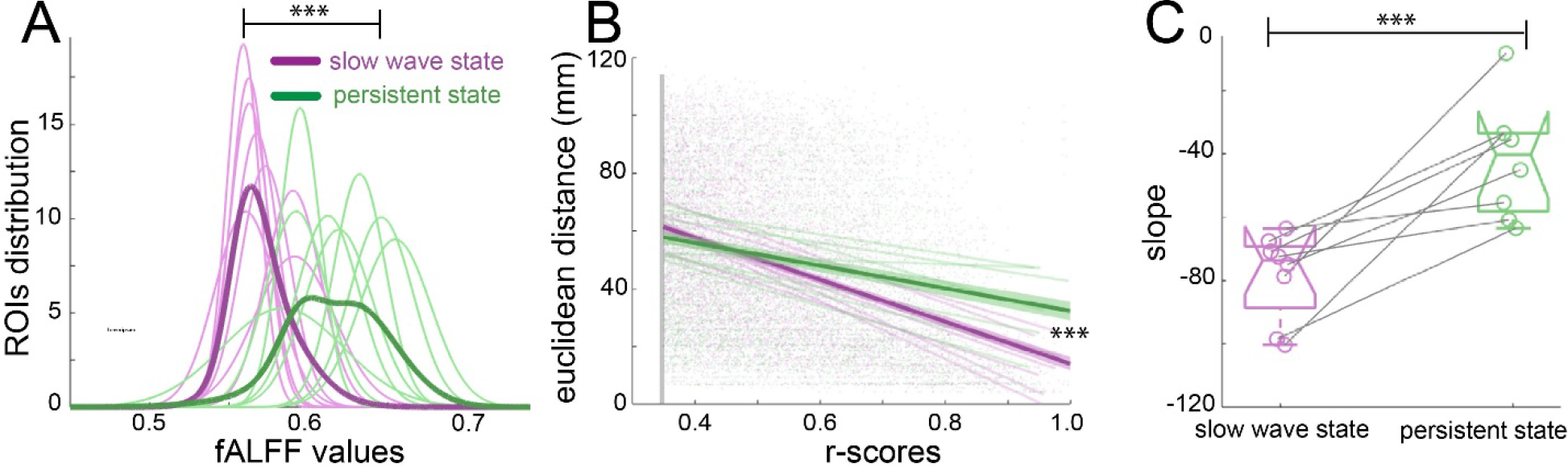
Network dynamics differentiate slow wave state and persistent state. A- fALFF distributions of the 96 cortical ROIs for each individual during slow wave state (magenta) and persistent state (green), ***p<=0.001. B- Correlation between the r-scores of each pair of cortical ROIs and their Euclidean distance, ***p<=0.001. C- Box plot of the slopes for each individual plotted in B. ***p<=0.001.

### Graph theory analysis

To assess the organizational nature of the identified cortical networks, we applied graph theory methods^51^. We first computed the modularity degree of the cortical networks during persistent state and slow wave state. Modularity is a quality index for a partition of a network into non-overlapping communities^54^. Therefore, in a brain with more compartmentalized networks this measure is higher than in one that shows a less selective network configuration. When we compared modularity values from networks during slow wave state with those from networks during persistent state we indeed observed significantly larger values in persistent state (t(7)=3.68, p=0.007; Figure 4A). Two important characteristics to differentiate small world from random networks are global and local efficiency. Small world networks exhibit high values for both measurements whilst randomized networks do show high values in global but low values for local efficiency^55^. When we compared these measures, we did not find significant differences in the global efficiency scores (**Figure 4B**) (t(7)=0.902, p=0.397) but we found significantly lower scores for slow wave state than for persistent state in the local efficiency measurement (**Figure 4C**) (t(7)=4.393, p=0.003). It is well documented that both, small world and random networks show short characteristic path length, whilst only small world networks show a high clustering coefficient^56^. In our analyses we found a significantly lower clustering coefficient for slow wave state (**Figure 4D**) (t(7)=3.323, p=0.011), but no significant differences between slow wave state and persistent state for the characteristic path length (**Figure 4E**) (t(7)=0.280, p=0.787), suggesting the idea that the cortex during slow wave state behaves like a random network, whilst during persistent state it behaves as small world network (**Figure 4F**). In summary, graph theory analysis indicates a different functional connectivity patterns in persistent state and slow wave state. In the first case it represents canonical well compartmentalized, small world networks, whilst in the second, a massively interconnected randomized network.

**Figure 4:**
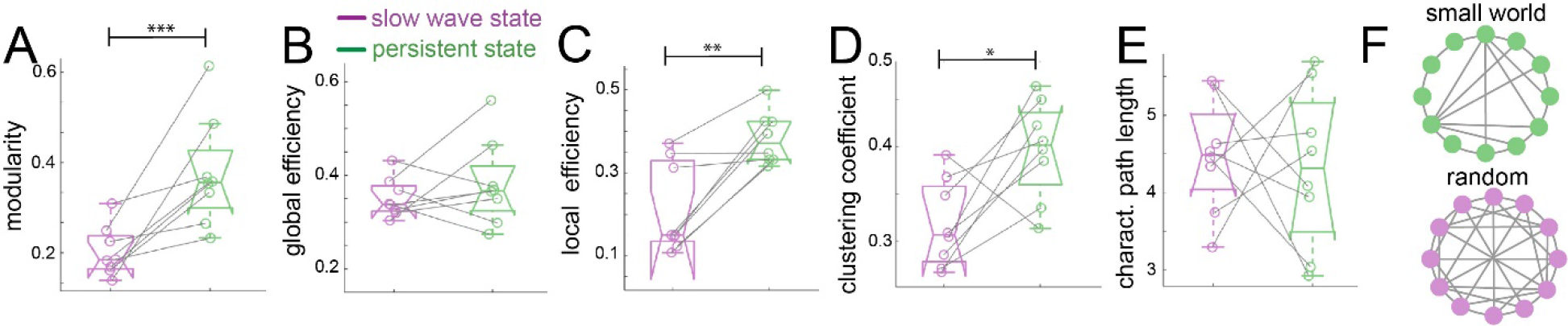
Graph theory shows a random network signature in slow wave state. *=p<0.05; **=p<0.01; ***=p<0.001; slow wave state (magenta), persistent state (green). A- modularity values. B- global efficiency values. C- local efficiency values. D- clustering coefficients. E- characteristic path lengths. F- Diagram representing a characteristic configuration of small world (green) and randomized networks (magenta).

### Connectivity differences are reflecting different brain states

Functional connectivity patterns in the rodent brain can change dramatically depending on the anesthetics used^13,57,58^. In order to rule out anesthesia-specific effects in our brain state classification, we applied a low dose of isoflurane condition (0.5%-0.7%) (n = 6 animals). In the raw data obtained by optic-fiber based calcium recordings (**Figure 5A**) it becomes apparent that the signals recorded under low isoflurane anesthesia resemble the ones obtained under medetomidine rather than the ones recorded under high isoflurane concentrations (**Figure 5B**). To quantify whether this observation is meaningful for the BOLD signal, we computed the functional connectivity analysis in the same way as for the previous experiments, to obtain the connectivity matrices (**Figure 5C**). We computed matrix similarity analysis for each of the 6 matrices against the mean matrix for slow wave state and persistent state to determine if the connectivity patterns obtained under low concentration of isoflurane resemble more the ones previously measured under medetomidine (persistent-state) or the ones measured under higher isoflurane concentrations (slow wave state). Similarity values can be compared extracting the eigen values of the matrix and then using *Frobenius norm* in order to calculate the similarity. The lower the obtained value is, the greater the similarity between two matrices. In this case, the resulting values of persistent state induced by low isoflurane, were significantly lower when compared with the ones for persistent state induced by medetomidine sedation than slow wave state under high isoflurane concentration (**Figure 4D**) (t(5)=5.19, p=0.003). We used the same matrix similarity computation in a dynamic connectivity analysis. Therefore, we compared the initial 5 minutes of measurement with a moving average of 5 minutes in steps of 1 minute. The results show that in the case of persistent state induced by medetomidine (PS-M) and by low isoflurane concentrations (PS-I) network configurations are significantly more stable than in the case of slow wave state induced by high isoflurane concentrations (**Figure 4E**). This result indicates that the network dynamics during SWS is related to a constant reorganization of connectivity patterns, probably due to the frequent changes in the rate of population *down-up transitions*.

**Figure 5:**
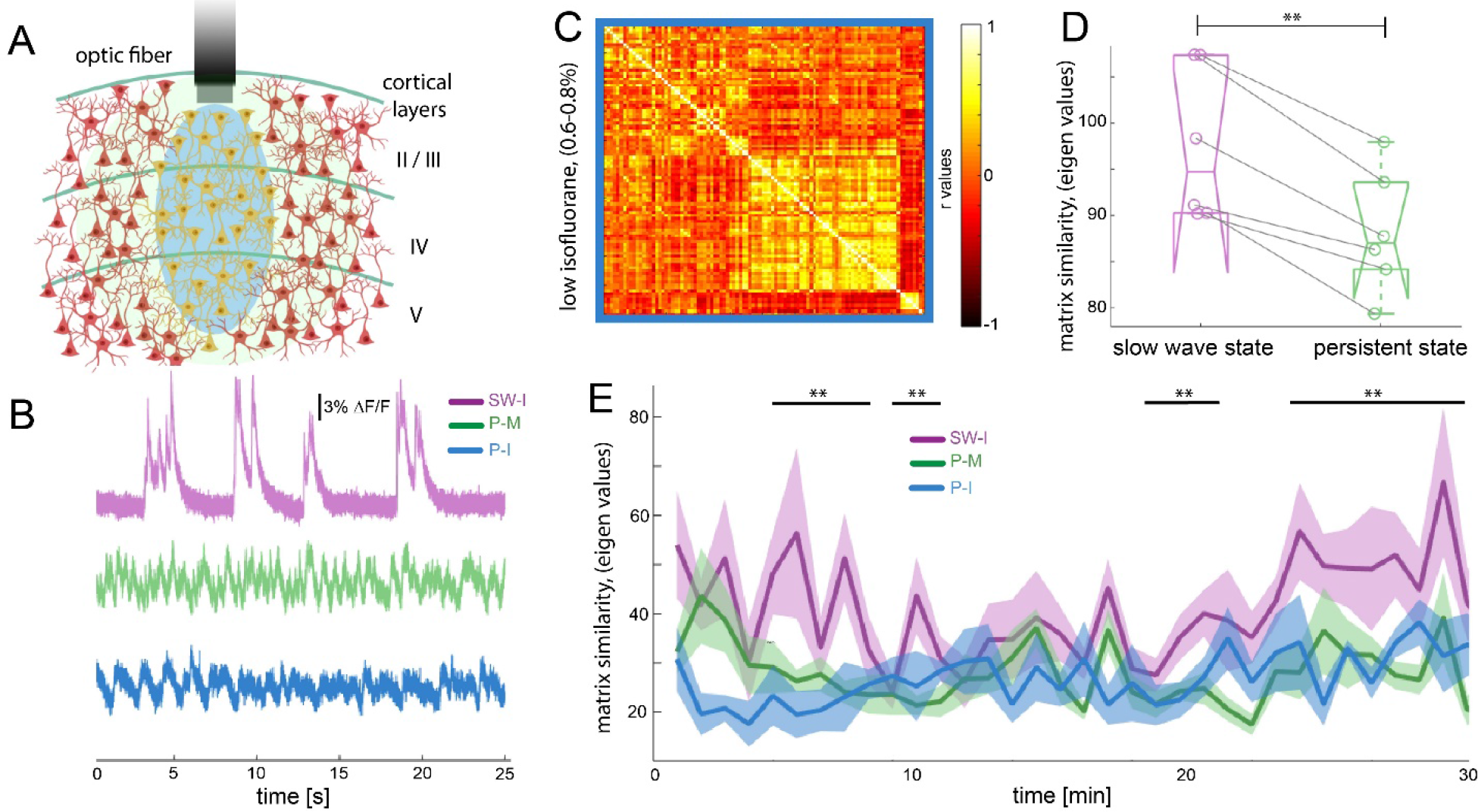
Different brain states result in characteristic connectivity matrices. A- Diagram of the optic-fiber based calcium recordings. OGB-1 was Bolus-injected, leading to a stained region with a circular geometry and a diameter of about 800 μm, depicted in light green. The optic fiber with a diameter of 200 μm was implanted at a cortical depth of about 300 μm, and blue light used for the excitation of the indicator, depicted in blue. Emitted light, comprising the changes in cytosolic calcium concentration of the local neural population was collected by the same fiber, and recorded by the fiber optometer. B- Characteristic traces of fiber recordings in slow wave state (SW-I, magenta), and persistent state induced by medetomidine (P-M, green) and low concentration of isoflurane (blue, P-I). C- Average of the correlation matrices (n=6) of the mean BOLD signal in 96 cortical regions under low isoflurane concentration. D- Matrix similarity analysis of the low isoflurane (P-I n=6) experiments compared with the average matrix generated under slow wave activity (SW-state, magenta) and persistent activity (P-M, green), **p<0.01. E- matrix similarity observed under the three conditions under a dynamic connectivity analysis. The matrices generated in a 5 minutes period with steps of 1 minutes where compared with the average matrix generated in the first 5 minutes for each individual, **p<0.01.

### Population down-up transitions drive functional connectivity during slow wave state

Slow wave state has been proposed to constitute the default activity pattern of the cortex ^21^; This bimodal activity pattern – periods of quiescence interrupted by short bursts of activity - generates waves that can propagate across the cortex ^24,59^. We recently showed that these waves generate a cortex-wide increase in BOLD activity^11^ which may be responsible for the cortex-wide increase in functional connectivity in slow wave state. To test this, we first reanalyzed the connectivity results during slow wave state using the cortex-wide component (CWC) as a covariable, since this component is a direct local readout of population slow wave state^11^. Therefore, if the high connectivity signature found during slow wave state is related to slow wave *down-up* transitions, the cortex-wide component as a covariable should remove any correlation that occurs due to these *down-up* transitions. Following this rationale, we expected that once the *down-up* transition covariable is removed from the connectivity analysis, correlation values should significantly decrease (**Figure 6Ai**). When comparing the cumulative distribution function of the r-values obtained using the inverted signal of the CWC (CWCinv) as a covariable, with those obtained using the original CWC as a covariable (CWCcov). The CDF (same analysis than Figure 2B) data showed significantly higher values for the -cwc condition at the mean of the cutoff (**Figure 6Aii**) (t(7)=4.756, p<=0.001). Furthermore, the amount of significantly correlated pairs for each individual at its particular cutoff was also significantly less in the CWCcov condition. (t(7)=3.742, p=0.007).This suggests a relationship between slow wave state-related *down-up* transitions and the strength of cortical functional connectivity. In order to obtain direct evidence, we co-registered optic-fiber-based calcium recordings ^60,61^ and fMRI BOLD during slow wave state in 5 animals (**Figure 6B-I**). We quantified the number of population *down-up* transitions in a period of 3 minutes in the calcium signal emitted by a local neural population in visual cortex (V1) (**Figure 6B-II, upper panel**) and correlated it with the number of significant pairs form the functional connectivity analysis of the BOLD signal for the same time period (**Figure 6B-I, lower panel**). We quantified these values for 30-minute-long recordings using samples of 3 minutes and a gap of 1 minute between them. The correlation between the amount of *down-up* transitions in the calcium signal and the significant pairs was highly significant in 4 of the 5 animals (p<0.001) (**Figure 6B-II**). These results suggest that slow wave state-related *down-up* transitions are directly linked to the increase in functional connectivity in slow wave state.

**Figure 6:**
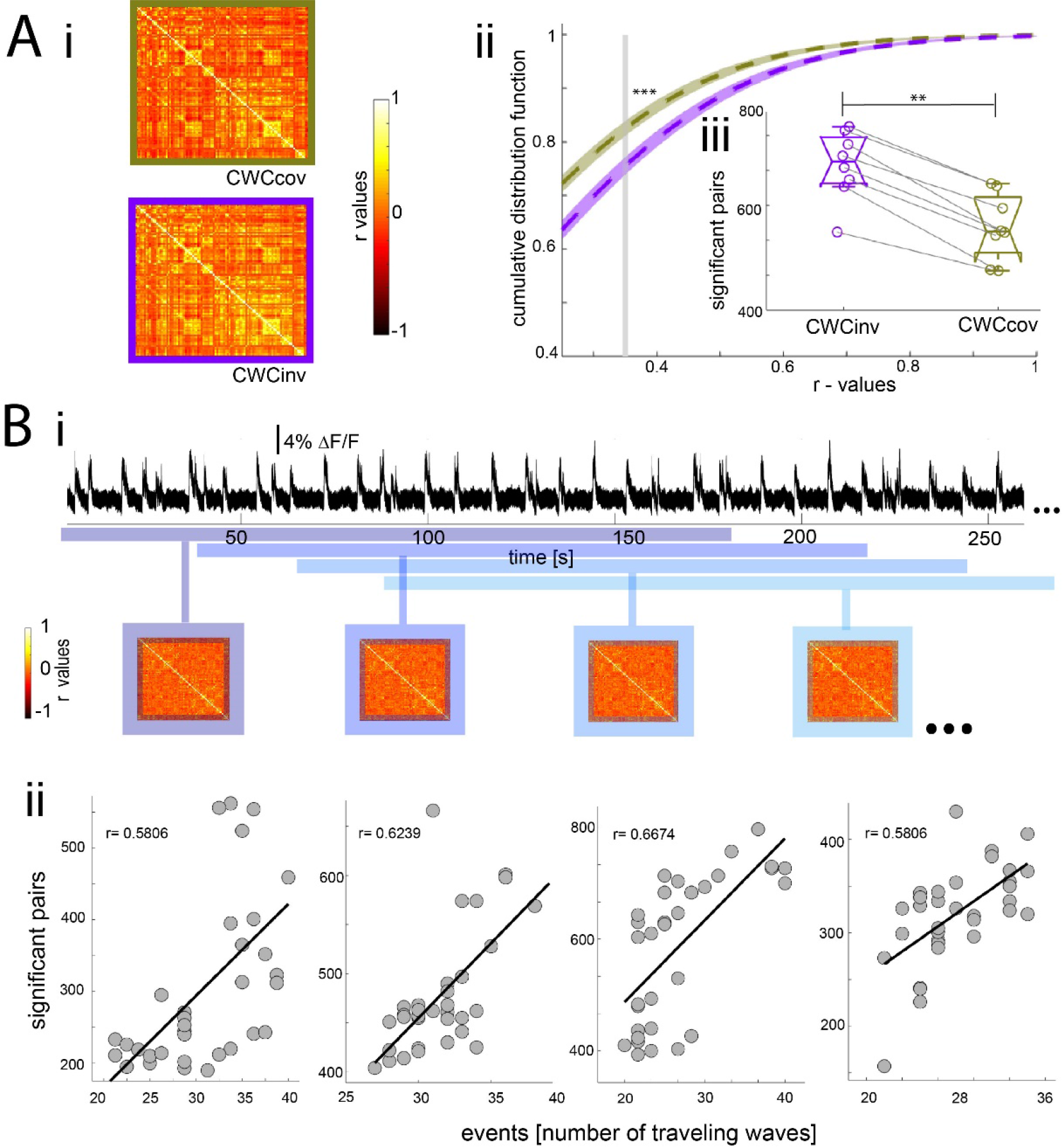
Population down-up transitions drive functional connectivity in slow wave state. A- functional connectivity of the BOLD signal using the cortex wide component of the ICA as covariable(CWCcov): i- Average of the correlation matrices (n=8) of the mean BOLD signal in 96 cortical regions under slow wave activity using the pan-cortical component of the ICA as covariable (CWCcov, purple framed) or the flipped values of the same component as control (CWCinv, brown framed). ii- Cumulative distribution curve for the r-values of the correlations under slow wave state or using the CWC of the ICA as regressor (CWCcov) or the flipped values of the same component (CWCinv). The vertical gray line points the mean of the FDR values used as cutoff to identify significant correlations, ***p<0.001. iii- Plot of the number of significant correlations for each individual in both conditions **p<0.01. B- Correlation between the amount significant pairs in the functional connectivity matrix and the number of down-up transitions. i- Correlation scheme. The number of down-up transitions was quantified for the respective time period (upper panel) and then correlated with the number of significant pairs obtained from the functional connectivity matrix (lower panel). ii- 4 of the 5 animals analyzed showed a significant positive correlation between number of down-up transitions and number of significant p functionally connected pairs.

### Simultaneous optic-fiber based calcium recordings in two cortical regions

While these previous results demonstrated the role of *down-up* transitions in functional connectivity during slow wave state, they are based on the BOLD activity and not on a direct measure of neural activity. Slow wave state of a spatially confined neural population can be detected by optic-fiber-based calcium recordings ^60,61^. To corroborate our results at the neural level, we conducted experiments employing simultaneous optic-fiber-based calcium recordings in somatosensory (S1) and visual (V1) cortex (**Figure 7A**). During slow wave state *down-up* transitions can be easily identified and quantified^23^. These transitions have a duration of 2 seconds on average (**Figure 7C**). The amplitude of the cross-correlation between these two signals during slow wave state is highly significant (**Figure 7D**) with a delay matching the propagation time of the wave moving from V1 to S1. To demonstrate that *down-up* transitions are underlying the increase in cortical connectivity found in our slow wave state data, we filtered the calcium and the BOLD signal at different frequency intervals and again run the cross-correlation analysis (**Figure 7D**). Results of 6 recordings in 3 animals show that the cross-correlation peaks significantly decrease in the respective frequency interval in which the *down-up* transitions are filtered out (**Figure 7E**). The cross-correlation values were significantly lower when data was filtered between 0.01-0.4 hz (t(5)=4.26, p=0.008), 0.2-0.6 hz (t(5)=7.99, p<0.001), and 0.3-0.7 (t(5)=7.28, p<0.001) hz. These results support those obtained previously with the BOLD connectivity analyses at the cortical population level, demonstrating *down-up* transitions to be the main contributor to the high index of cortical connectivity found during slow wave state.

**Figure 7:**
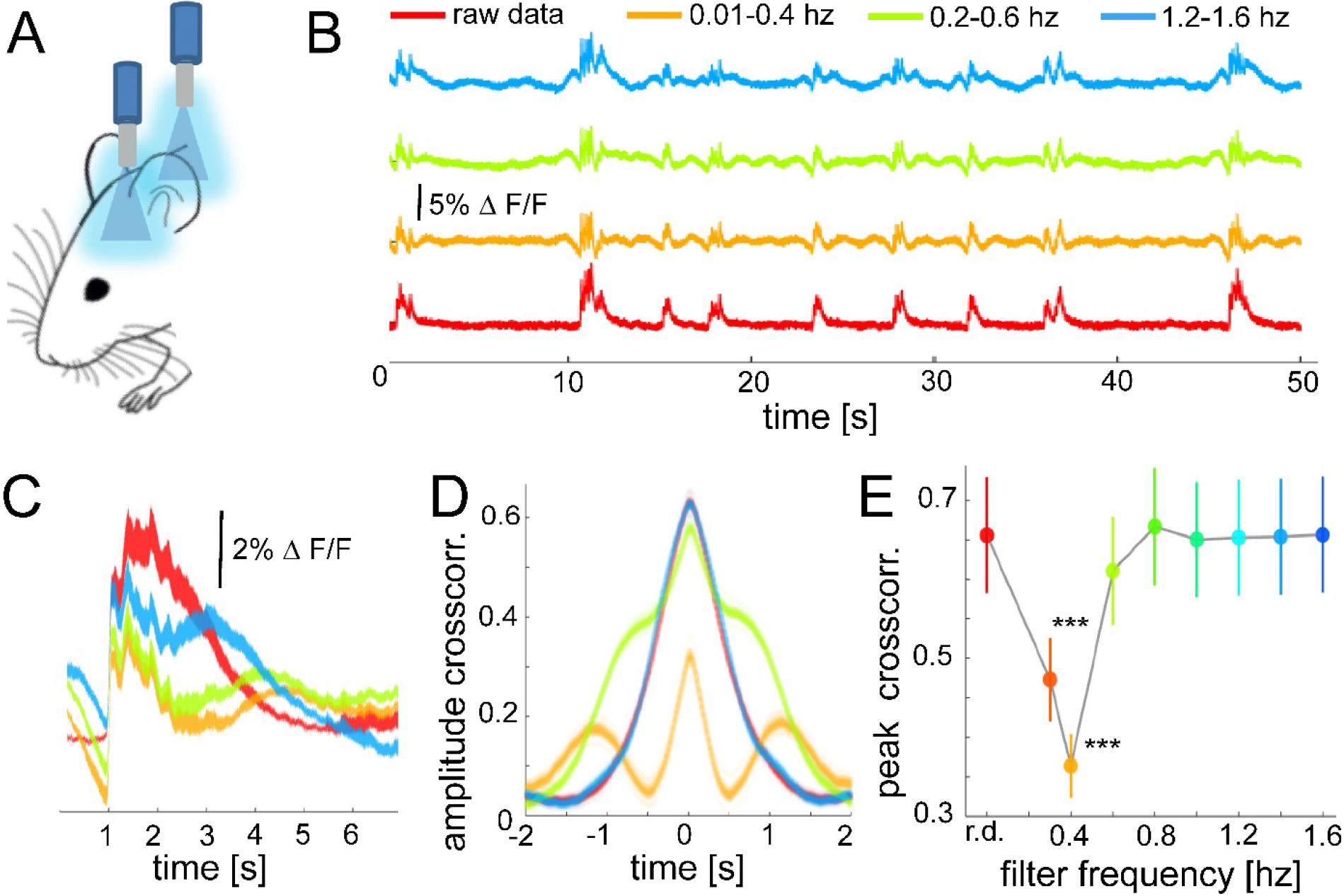
In slow wave state, the correlation between distant optical recordings is driven by slow-wave-associated calcium waves. A- Diagram of dual-simultaneous optic-fiber based calcium recordings B- Exemplary calcium trace filtered at different band-stop intervals, excluding those particular intervals from the recordings. C- Mean of the down-up transitions for a trace (◻f/f) at different band stop filters. D- Cross-correlation results of a single recording at different band-stop intervals. E- Average of the cross-correlation peak values (n=6) at different band-stop intervals, ***p<0.001.

## Discussion

In this study, we demonstrated that brain states can modify properties of the functional connectome through changes in cortical excitability. Independent component analysis showed distinct functional networks for each of the investigated states. In the case of slow wave state a cortex-wide component was found in all the animals, while in persistent state, a clear compartmentalization of the cortical structures gave origin to well determined networks, as the default mode or the auditory network. Notably, the same brain state could be found using two different types of anesthesia, suggesting, that not the type of used anesthesia but the brain state induced by the level of anesthesia influences resting state networks. The functional connectivity analyses using cortical parcellation corroborated this concept, showing an increase in the connectivity r-scores as in the number of significant correlations in slow wave state. Networks during slow wave activity are characterized by long periods of quiescence followed by short lapses of activity corresponding to a propagating wave^24^, as demonstrated by the simultaneous cortical calcium recordings. To rule out that the connectivity values observed during slow wave activity are due to the spurious consequence of an increase in power we performed fALFF analysis and found significantly lower values in the slow wave state, probably due to the long periods of neuronal quiescence that characterize the slow wave state.

### Spatiotemporal network dynamics

The high degree of compartmentalization in persistent state observed in the ICA can be corroborated by calculating the dependency of the distance in the r-scores between two areas. In a highly compartmentalized connectome, this dependency should be lower than in a system which is highly interconnected ^62^. Indeed, when correlating the r-scores and the Euclidean distance of the network’s pairs for persistent state, the correlation was significantly lower than for slow wave state. This finding suggests, that indeed, the functional architecture in persistent state is dominated by specific compartmentalized network nodes, which are not necessarily spatially close, such as the default mode network. In sharp contrast, in slow wave state, we observed a strong distance dependency, in line with a travelling wave of excitability – a slow wave event – eventually recruiting the entire cortex. These travelling slow waves temporarily switch the functional network from hyperpolarized rather quiescent down state into a pan-cortical rather depolarized up state of high excitability ^21,22,63^.

Applying graph theory methods, we found that during persistent state the network shows characteristic small world network dynamics, as they have been described for the awake human brain ^35^. During slow wave state graph theory analysis showed characteristics of highly interconnected random networks, corroborating the idea of a lack of a spatially segregated functional connectivity signature in that state ^36,64^.

One could argue that the difference in connectivity is reflecting a particular change in the neural dynamics induced by the anesthetic regimen rather than presenting a correlate of a particular state of the brain, since it has been reported that functional connectivity patterns in the rodent brain change dramatically depending on the anesthetics used ^13,57,58^. In order to rule out anesthesia-specific effects on connectivity we demonstrated that decreasing the amount of isoflurane administrated to the animal let to the transition of slow wave state to persistent state. And indeed, the functional connectivity dynamics observed under a low dose of isoflurane resembled the pattern observed in persistent state, rather than the one observed in slow wave state.

Recently is has been shown that spontaneous task-free brain activity is non-random in the temporal domain, and that this feature is well conserved between rodents and humans ^19^. Interestingly, when we analyze the temporal dynamics of connectivity patterns, we found that animals in persistent state, induced either by medetomidine or a low dose of isoflurane, showed rather temporally stable connectivity dynamics, corroborating the notion of an indeed persistent state ^22,28,63^. In contrast, animals in slow wave state showed fast, and temporally irregular transitions. This suggests, that individual, specific events within the slow wave state drive the state-specific network signature. The occurrence of these events, down-up transitions, or slow waves, are not occurring in a regular, oscillation-like manner, but depend on the overall physiological state of the animal, such as the depth of anesthesia ^5^. Although, there is abundant evidence on the relevance of infra-slow electrical brain activity for resting state connectivity^34,65,66^, there has been no direct neural evidence so far linking brain states to changes in network configuration.

When we used the cortex-wide component as a covariable in the connectivity analyses, the connectivity values decreased significantly. This component is a direct local readout of population slow wave state^11^ and is therefore representing the occurrence rate of down-up transitions. Furthermore, a direct demonstration of the involvement of down-up transitions in the functional connectivity maps comes from the significant correlation between the degree of connectivity and the amount of transitions in 4 of 5 animals.

To provide direct evidence for this hypothesis, we conducted simultaneous optic-fiber based calcium recordings alongside fMRI measurements in slow wave state, as previously implemented ^67^. When the component of down up transitions identified in the optical neural recordings were filtered out, we indeed found a decrease in correlation of cortical activity. These findings provide additional evidence, that indeed these distinct transitions in excitability drive the cortical functional connectivity in this brain state.

It is widely known that the generation and propagation of slow waves is depending on cortical excitability during sleep ^68^ or under anesthesia ^69^, even differentially affecting cortical layers ^70^. Slow wave propagation can even be reflective of a pathophysiological condition such as early AD ^71^. We propose, that distinct brain states, i.e. defined spatio-temporal dynamics of cortical excitability, drive the functional connectivity of the brain. These rather discrete states – persistent vs slow wave state, are characterized by particular and well-defined network signatures. This new understanding of the network dynamics underlying the respective functional network pattern will reduce the variability and increase translational valence of task-free resting state study

### Implications on resting state studies

Several studies have demonstrated that the functional state of the brain is dynamic over time, and this is reflected in functional connectivity changes ^72,73^. These dynamic states are quantifiable, depend on the ongoing infraslow activity and seem to show atypical dynamics in some neurological disorders as autism ^74^. Degrees of wakefulness seem to play an important role in global functional connectivity ^75^. For instance, Tagliazucchi & Laufs showed that during resting state recordings in humans the dynamic changes in the functional neuroanatomy are due to switches in wakefulness and sleep patterns^17^. More recently Laumann and colleagues showed that the dynamic variability over time in the fMRI functional connectivity signatures can be explained by fluctuating sleep stages^76^.

Previous task-free fMRI studies report that anesthetized animals show lower functional coupling than awake animals ^77,78^. However, some studies could show that default mode networks in rodents can be found even after the loss of consciousness ^31,79^, while others show a breakdown in the networks of resting state connectivity under anesthesia ^80,81^. These apparent dissimilar results have been hypothesized to be dependent on the synchronization of infra-slow oscillatory activity and cross-frequency coupling^82^. Our experiments demonstrate that the origin of these discrepancies rely in the different brain states. During persistent state compartmentalized activity is detected, whilst in slow wave state the degree of connectivity depends in the rate of slow waves generated in a particular period of time. Therefore, these differing results in the network configurations can be merely due to different brain states or under slow wave, the amount of down-up transitions. Brain states and the rate of down-up transitions are dynamic, and both states can be found under the same conditions, e.g. during wakefulness ^83^. Here, we present data from strictly controlled conditions, in which the entire cortex seems to be engaged in either persistent state or slow wave state, however, in awake or slow wave sleep, this might not be the case. This calls for a brain-state-informed analysis of functional connectivity in task-free “resting state” studies, both in humans as well as in animal models. Human subjects may transition between these brain states when they fall asleep ^84^. Even more so, animals react differently to the same anaesthetic regimen ^83^. It might be the case, that at the beginning of an imaging experiment, an animal is in persistent state, and further along the anaesthesia effect increases, shifting the animal into slow wave state, with drastically different connectivity signatures. Therefore, it is necessary to consider the particular brain state and global changes in cortical excitability before driving conclusions about networks dynamics in task-free connectivity analyses. It might be advisable, to probe the brain state before and after a connectivity experiment. This might allow us to ameliorate the translational roadblock by removing the variance of different brain states relating human and rodent-task free connectivity analyses.

## Acknowledgments

This work was supported by the DFG (SPP 1665: Resolving and manipulating neuronal networks in the mammalian brain - from correlative to causal analysis; CRC1193: Neurobiology of resilience to stress-related mental dysfunction: from understanding mechanisms to promoting preventions) and the Focus Program translational Neurosciences (FTN-Mainz). Authors would like to thank Nuse Afahaene for excellent technical assistance and Eduardo Rosales Jubal for support in the Graph theory analyses.

